# Chondroitinase and antidepressants promote plasticity by releasing TRKB from dephosphorylating control of PTPσ in parvalbumin neurons

**DOI:** 10.1101/2020.08.13.249615

**Authors:** Angelina Lesnikova, Plinio Cabrera Casarotto, Senem Merve Fred, Mikko Voipio, Frederike Winkel, Anna Steinzeig, Hanna Antila, Juzoh Umemori, Caroline Biojone, Eero Castrén

## Abstract

Perineuronal nets (PNNs) are an extracellular matrix structure rich in chondroitin sulphate proteoglycans (CSPGs) which preferentially encase parvalbumin-containing (PV+) interneurons. PNNs restrict cortical network plasticity but the molecular mechanisms involved are unclear. We found that reactivation of ocular dominance plasticity in the adult visual cortex induced by chondroitinase (chABC)-mediated PNN removal requires intact signaling by the neurotrophin receptor TRKB in PV+ neurons. Additionally, we demonstrate that chABC increases TRKB phosphorylation (pTRKB), while PNN component aggrecan attenuates BDNF-induced pTRKB in cortical neurons in culture. We further found that protein tyrosine phosphatase sigma (PTPσ, PTPRS), receptor for CSPGs, interacts with TRKB and restricts TRKB phosphorylation. PTPσ deletion increases phosphorylation of TRKB *in vitro* and *in vivo* in male and female mice, and juvenile-like plasticity is retained in the visual cortex of adult PTPσ deficient mice (PTPσ+/-). The antidepressant drug fluoxetine, which is known to promote TRKB phosphorylation and reopen critical period-like plasticity in the adult brain, disrupts the interaction between TRKB and PTPσ by binding to the transmembrane domain of TRKB. We propose that both chABC and fluoxetine reopen critical period-like plasticity in the adult visual cortex by promoting TRKB signaling in PV+ neurons through inhibition of TRKB dephosphorylation by the PTPσ-CSPG complex.

**Significance statement:** Critical period-like plasticity can be reactivated in the adult visual cortex through disruption of perineuronal nets (PNNs) by chondroitinase treatment, or by chronic antidepressant treatment. We now show that the effects of both chondroitinase and fluoxetine are mediated by the neurotrophin receptor TRKB in parvalbumin-containing (PV^+^) interneurons. We found that chondroitinase-induced visual cortical plasticity is dependent on TRKB in PV^+^ neurons. Protein tyrosine phosphatase type S (PTPσ, PTPRS), a receptor for PNNs, interacts with TRKB and inhibits its phosphorylation, and chondroitinase treatment or deletion of PTPσ increases TRKB phosphorylation. Antidepressant fluoxetine disrupts the interaction between TRKB and PTPσ, thereby increasing TRKB phosphorylation. Thus, juvenile-like plasticity induced by both chondroitinase and antidepressant treatment is mediated by TRKB activation in PV^+^ interneurons.

## Introduction

Plasticity is the ability of the brain to change itself through establishing new neuronal connections and rewiring existing ones. Plasticity is prominent in early life and it peaks during so-called “critical periods” when the ability of the brain to adapt is at its highest (Wiesel, 1982). After the end of the critical period, plasticity persists but at significantly diminished levels (Hübener and Bonhoeffer, 2014).

Closure of the critical periods is mediated by changes in cortical excitatory/inhibitory (E/I) balance that take place due to maturation of cortical inhibitory interneurons. Fast-spiking interneurons expressing parvalbumin (PV^+^) orchestrate synchronous neuronal oscillations and play a particularly important role in this process (Hensch, 2005). Closure of the critical period coincides with the functional maturation of PV-positive cells and establishment of perineuronal nets (PNNs) around them (Wang and Fawcett, 2012; Umemori et al., 2018; Fawcett et al., 2019). Perineuronal nets are mesh-like structures of extracellular matrix that surround the somata and proximal dendrites of PV^+^ interneurons in particular (Kwok et al., 2011). Chondroitin sulfate proteoglycans (CSPGs), such as aggrecan and brevican, are major components of PNNs.

After the closure of a critical period, neuronal plasticity can still be modulated, and critical period-like plasticity can be induced in the adult brain by a number of different methods (Bavelier et al., 2010; Sale et al., 2014). Digestion of perineuronal nets by chondroitinase (chABC) treatment has been demonstrated to induce ocular dominance plasticity in the adult visual cortex (Pizzorusso et al., 2002). Local chondroitinase injections into different brain areas have also been shown to promote recovery of spinal cord injury (Bradbury et al., 2002) and extinction of fear memories in adult rodents (Gogolla et al., 2009; Hylin et al., 2013; Shi et al., 2019). However, the mechanisms through which chABC influences plasticity in the central nervous system (CNS) remain unclear.

Antidepressant drugs also induce critical period-like plasticity in the adult brain (Castreń, 2013). Antidepressants activate neurotrophic receptor tyrosine kinase 2 (TRKB), the receptor for brain-derived neurotrophic factor (BDNF), and promote plasticity through its signaling pathways (Saarelainen et al., 2003; Duman and Monteggia, 2006; Castrén and Antila, 2017; Umemori et al., 2018). Like ChABC, fluoxetine, a widely prescribed antidepressant, induces ocular dominance plasticity in the rodent visual cortex (Maya Vetencourt et al., 2008) and makes fear-related memories in mice susceptible to erasure (Karpova et al., 2011). We have recently found that the activation of TRKB in PV^+^ interneurons is both necessary and sufficient for antidepressant-induced plasticity in the mature CNS (Winkel et al., 2020).

ChABC and antidepressant treatment exert similar plasticity-promoting effects in the adult brain; however, it is not known whether they recruit similar molecular mechanisms. We hypothesize that receptor-like protein tyrosine phosphatase sigma (PTPσ) might be a nexus that mediates plasticity processes induced by both methods. PTPσ is a receptor for chondroitin sulfate proteoglycans (CSPGs) (Shen et al., 2009), and it has been demonstrated to be essential for the inhibitory effects of CSPGs on neurite outgrowth (Shen et al., 2009; Duan and Giger, 2010; Coles et al., 2011). PTPσ interacts with and modulates the activity of TRK receptors (Faux et al., 2007; Takahashi et al., 2011), inhibiting TRKB through dephosphorylation (Kurihara and Yamashita, 2012).

We now demonstrate that TRKB activity in PV^+^ interneurons is essential for plasticity induced by chABC. While CSPGs removal by chABC injection into the visual cortex promotes ocular dominance shift, this effect is abolished in heterozygous mice with reduced TRKB in parvalbumin-positive neurons (PV-TRKB^+/-^). We have confirmed that PTPσ interacts with TRKB, and genetic deficiency of PTPσ promotes TRKB phosphorylation (pTRKB) *in vitro* and *in vivo.* We further show that PTPσ deficiency promotes plasticity at network levels, as PTPσ^+/-^ mice display critical period-like plasticity in the visual cortex in adulthood. Finally, we have observed that the antidepressant fluoxetine disrupts TRKB: PTPσ interaction *in vitro* and *in vivo*.

## Materials and Methods

### Animals

Balb/c and C57BL/6J mice heterozygous for *PTPRS* gene (PTPσ^+/-^ mice) and their wild-type littermates, C57BL/6J mice heterozygous for *TRKB* gene in parvalbumin-expressing (PV+)-interneurons (PV-TRKB^+/-^) and their PV-Cre littermates (PV-TRKB^+/+^), wild-type C57BL/6J mice were used in the experiments. Balb/c PTPσ^+/-^ mice were originally developed by Michel Tremblay’s lab (McGill University, Canada) (Elchebly et al., 1999) and kindly donated to us by Heikki Rauvala (University of Helsinki, Finland). Balb/c PTPσ^+/-^ mice 10 weeks old were used for brain sample collection and pTRKB level assessment by ELISA. For optical imaging experiments, balb/c mouse line was rederived to C57BL/6J background, and N4 generation of the offspring was used for testing. The mice were 2 months old at the beginning of the experiments. PV-TRKB^+/-^ (TRKB^flx/wt^, PV^cre/wt^) mice were generated by mating heterozygous floxed TRKB mice (TRKB^flx/wt^) (Minichiello et al., 1999) and homozygous PV-specific Cre line (PV^cre/cre^) (Pvalb-IRES-Cre, JAX: 008069, Jackson laboratory) (Hippenmeyer et al., 2005). PV heterozygous (TRKB^wt/wt^, PV^cre/wt^) littermates with intact TRKB in PV^+^ neurons were used as a control group for PV-TRKB^+/-^ mice. The mice were 4 months old at the beginning of the experiments. Wild-type C57BL/6J mice 5 months old were used to assess fluoxetine effect on TRKB:PTPσ interaction in the visual cortex *in vivo.* The mice were kept under standard laboratory conditions with 12-hour light/dark cycle (lights on at 6:00 am) and access to food and water *ad libitum.* All the procedures involving animals were done in compliance with the National Institutes of Health Guide for the Care and Use of Laboratory Animals guidelines and were approved by the Experimental Animal Ethical Committee of Southern Finland (ESAVI/10300/04.10.07/2016).

### Brain sample collection and processing

Mice were sacrificed with CO2. The death was confirmed by ascertaining cardiac and respiratory arrest. The animals were decapitated, the visual cortices were dissected and stored at −80° C. Samples from the primary visual cortex were sonicated in NP lysis buffer (137mM NaCl, 20mM Tris, 1% NP-40, 10% glycerol, 48mM NaF) containing a protease and phosphatase inhibitor mix (#P2714 and #P0044, Sigma Aldrich, USA) and 2mM Na2VO3. The homogenate was centrifuged for 15 min. at 15 000 G, 4° C. The supernatant was collected and used for further analysis. Protein levels were measured using DC Protein Assay Kit (Bio-Rad, USA, #5000116) by colorimetric Lowry method in Varioskan Flash (Thermo Fisher Scientific, USA).

### Cell culture

Cerebral cortical cell cultures were prepared from Wistar rat (Harlan labs, U.K) embryos extracted on embryonic day 18 (E18) (Sahu et al., 2019). The cells were cultured in a serum-free neurobasal medium with supplements (1% penicillin, 1% L-glutamine, 2% B-27%) and collected after 7-9 days in vitro (DIV). For the experiments assessing the effect of PTPσ genetic deficiency on pTRKB *in vitro,* cortical cell cultures were prepared from balb/c mouse embryos (PTPσΛ PTPσ^+/-^ and their WT littermates) extracted on embryonic day 18 (E18) using the same protocol as described above for the rat cells. Mouse fibroblast cells stably expressing full-length TRKB (MG87.TRKB) or TRKA (MG87.TRKA) were cultured in Dulbecco’s Modified Eagle’s Medium (DMEM) supplemented with 10% fetal calf serum (FCS), 1% penicillin/streptomycin (PS), 1% L-glutamine and 400 mg/ml G418. Human-derived HEK293T cells were cultured in DMEM supplemented with 10% FCS, 1% PS and 1% L-glutamine.

### Transfection

70% confluent MG87.TRKA, MG87.TRKB and HEK293T cell lines were transfected using Lipofectamine 2000 (Thermo Fischer Scientific, USA). MG87.TRKA and MG87.TRKB cells were transfected with PTPσ (RefSeq number NM_019140) Myc-DDK-tagged open-reading frame (ORF) plasmid purchased from OriGene (#RR209636), pCMV6-Entry vector with C-terminal Myc-DDK Tag (#PS100001) was used to transfect control cells. HEK293T were transfected with GFP-tagged full-length TRKB plasmid (HP220GFP-TRKB)(Haapasalo et al., 2001) or TRKB carrying Y433F/R427A mutation (Cannarozzo et al., 2020).

### Antibodies and reagents

Aggrecan was purchased from Sigma-Aldrich, USA (A1960). Anti-phospho-TRKA (Tyr490)/TRKB (Tyr516) (C35G9) rabbit monoclonal antibody (mAb) was purchased from Cell Signaling Technology, USA (#4619, #4621 and #4168). Anti-TRKB goat polyclonal antibody (pAb) was bought from R&D Systems, USA, #AF1494. Anti-phosphotyrosine mouse monoclonal antibody (clone PY20) was purchased from Bio-Rad, USA (#MCA2472). Anti-PTPσ (SS-8) mAb was purchased from Santa Cruz Biotechnology, USA (SC-100419). Secondary HRP-conjugated antibodies were purchased from Bio-Rad, USA (goat anti-rabbit #1705046 and goat anti-mouse #1705047) and from Invitrogen, USA (rabbit anti-goat #611620). Enhanced chemiluminescent (ECL) substrate WesternBright Quantum horseradish peroxidase (HRP) substrate (Advansta, USA) was used for Western blot (#K-12042). Pierce HRP ECL substrate was used to detect luminescence in ELISA (Thermo Fisher Scientific, USA, #32209). Tris-buffered saline with Tween-20 (TBST) was used for Western blot (20 mM Tris–HCl; 150 mM NaCl; 0.1% Tween-20; pH 7.6), phosphate-buffered saline with Tween-20 (PBST) was used for ELISA (137 mM NaCl; 10 mM Phosphate; 2.7 mM KCl; 0.1% Tween-20; pH 7.4).

### Drug treatment

10 μg/ml aggrecan was added to cortical neurons cultured 6 DIV, and the cells were collected 1 day after the aggrecan treatment (7 DIV). 20 ng/ml BDNF was added to cortical cells for 10 min. 0.1 and 1 μM fluoxetine was added to cortical cells cultured 7 DIV, and the cells were collected after 30 minutes of treatment. HEK293T cells were treated with 10 μM fluoxetine for 30 min. Balb/c PTPσ^+/-^ and wild type mice were injected with 30 mg/kg fluoxetine intraperitoneally and sacrificed with CO2. The brain samples were collected and stored at −80 C° until further processing.

### Immunoprecipitation and Western blot

The cells were lysated using NP lysis buffer containing 2 mM sodium orthovanadate and protease inhibitor mix. The homogenized suspension was centrifuged (15000 G, 10 min, +4 °C), and the resulting supernatant was used for analysis. For immunoprecipitation, TRKB was captured using anti-TRKB antibodies (R&D Systems, USA). The samples were incubated with sepharose, washed with NP lysis buffer twice and the proteins were separated by heating in 2X Laemmli buffer (4% SDS, 20% glycerol, 10% 2-mercaptoethanol, 0.02% bromophenol blue and 125mM Tris HCl, pH 6.8) for 5 min. at 95° C. The samples were loaded to NuPAGE 4-12% Bis-Tris Protein polyacrylamide gels (Invitrogen, USA, #NP0323BOX), and the proteins were separated according to their molecular weight using electrophoresis. The samples were transferred to polyvinylidene difluoride (PVDF) membrane, incubated in 1:1000 primary antibody dilution in 3% bovine serum albumin (BSA) in tris-phosphate buffer containing 0.1% Tween-20 (TBST) overnight at 4° C and subsequently incubated in horseradish peroxidase (HRP) conjugated secondary antibodies (1:10000) for 1 hour at room temperature (RT). The bands were visualized using chemiluminescent western blotting substrate in Fuji LAS3000 camera (Tamro Medlabs, Finland).

### ELISA

Levels of TRKB phosphorylation (pTRKB) were evaluated using enzyme-linked immunosorbent assay (ELISA) for pTRKB developed in our lab (Vesa et al. 2011). On day 1, the high-binding 96-well OptiPlate (Perkin Elmer, USA) was incubated overnight at 4° C with 1:500 anti-TRKB antibody diluted in a homemade carbonate buffer (57.4 mM sodium bicarbonate, 42.6 mM sodium carbonate, pH=9.8). On day 2, the plate was incubated for 2 hours at RT in the blocking buffer (3% BSA in PBST) to block non-specific binding. Homogenized and centrifuged brain samples or lysated cell samples were added to the plate and incubated overnight at 4° C. On day 3, the plate was washed 4 times with PBST using an automated plate washer (Thermo Fisher Scientific Wellwash Versa, USA), and the samples were incubated in anti-pTRKB antibodies diluted 1:1000 in the blocking buffer overnight at 4° C. On day 4, the samples were washed 4 times in PBST and incubated with tertiary HRP-conjugated antibodies in the blocking buffer (1:5000) at RT for 1 hour. Finally, ECL was added to the plate and luminescence was measured with 1 s integration time using Varioskan Flash plate reader (Thermo Fisher Scientific, USA). Levels of TRKB:PTPσ interaction were evaluated by ELISA using a similar protocol. TRKB was captured using anti-TRKB antibodies, and anti-PTPσ were applied afterwards to assess levels of PTPσ bound to TRKB.

### Cell-surface ELISA

Cell-surface ELISA was carried out to assess PTPσ levels found on the cell surface as previously described (Zheng et al., 2008; Fred et al., 2019). Briefly, cortical cells were cultivated in clear-bottom 96-well plates (ViewPlate 96, Perkin Elmer). On 7 DIV the cells’ medium was removed, they were washed with cold PBS and fixed with 100 μl of 4% paraformaldehyde (PFA) per well for 20 min. Then the cells were washed with PBS 3 times, and non-specific binding was blocked with PBS containing 5% non-fat dry milk and 5% BSA for 1 hour at RT. After that, the samples were incubated in primary anti-PTPσ antibody (1:500 in the blocking buffer) overnight at 4° C. On the following day, the cells were washed with PBS once and incubated in HRP-conjugated anti-mouse antibody (1:5000 in the blocking buffer) for 1 h at RT. The cells were washed 4 times with PBS, ECL was added and chemiluminescence was measured with 1 s integration time using Varioskan Flash (Thermo Fisher Scientific, USA).

### Transparent skull surgery

Transparent skull surgery was carried out as described in (Steinzeig et al., 2017). Animals were anesthetized either with a mixture of 0.05 mg/kg Fentanyl (Hameln, Germany), 5 mg/kg Midazolam (Hameln, Germany) and 0.5 mg/kg Medetomidine (Orion Pharma, Finland) administered intraperitoneally (i.p.). or isoflurane combined with 0.05 mg/kg buprenorphine analgesia administered subcutaneously (s.c.). 5 mg/kg carprofen (ScanVet, Nord Ireland) was administered s.c. for postoperative analgesia. Under anesthesia, the scalp and periosteum of the animals were removed, the skull was polished, and two layers of transparent acryl powder (EUBECOS, Germany) mixed with methyl methacrylate liquid (Dentsply, Germany) were applied on the surface. Metal holders (Neurotar, Finland) were installed on the top of the head and fixed with a mixture of acryl polymer powder (Dentsply, Germany) and cyanoacrylate glue (Steinzeig et al., 2017).

### Monocular deprivation

Monocular deprivation (MD) was carried out by suturing the eye contralateral to the imaged hemisphere (left eye) with perma-hand silk thread (Ethicon, USA). The length of the MD was 3.5 days for the PTPσ^+/-^ vs WT experiment and 7 days for the PV-TRKB^+/-^ vs WT experiment. The integrity of the suture was checked on a daily basis before the lights were on.

### Optical Imaging

Optical imaging of intrinsic signals was carried out as previously described in (Steinzeig et al., 2017). Visual cortex of the right hemisphere of each animal was imaged. Continuous-periodic stimulation with continuous synchronized data acquisition was used for the processing of the intrinsic signals. A drifting thin horizontal bar 2° wide moving upwards with a temporal frequency of 1 cycle/ 8.3 s (0.125 Hz) and a spatial frequency of 1/80° was utilized to alternatively stimulate the left and right eye, while the other eye was patched. The dirfting bars were displayed −15° to +5° from the center to optimally stimulate the binocular area of the right visual cortex. The imaging was done under 1.2% isoflurane anesthesia in a 1:2 mixture of O2:air.

### Optical imaging of PV-TRKB^+/-^ mice

Four-month-old mice were used for the experiments. During week 1, the animals underwent transparent skull surgery. After 7 days, during week 2, the animals underwent the first session of the optical imaging under isoflurane anesthesia (IOS1). During week 3, the animals were injected with 50 mU of chABC in PBS or PBS into the binocular area of the visual cortex and were subjected to monocular deprivation (MD) for 7 days. During week 4, the eyes were opened, and IOS2 immediately took place.

### Optical imaging of PTPα^+/-^ mice

Two-month-old mice were used for experiments. During week 1, the animals underwent transparent skull surgery. After 7 days, the animals underwent the first session of the optical imaging (IOS1) followed by MD for 3.5 days. We have switched to the shorter monocular deprivation length as compared to the previously described experiment since it has been recently demonstrated to be sufficient for the induction of the critical period-like plasticity (Baho et al., 2019). On day 4 of the monocular deprivation, the eyes were opened, and IOS2 took place.

### Stereotaxic surgeries

The mice were anesthetized with isoflurane combined with 0.05 mg/kg buprenorphine analgesia. Images acquired during the first session of the optical imaging were used to identify the binocular area of the visual cortex. A hole in the skull was made with a drill, and 50 mU of chABC in 1 μl of PBS or 1 μl of PBS were injected in the center of the binocular area using a microsyringe pump. 10 μl Nanofil syringe (WPI Nanofil) with drilled 1.1 mm outlet bore to accommodate 1.0 mm glass needle and custom-made bevelled borosilicate glass needles with a 50 μm tip diameter were used for the infusions. Infusions were done at the speed of 2 nl/sec. After the surgery, the animals were left to recover in the home cage, and the second imaging session took place 7 days later.

### Experimental design and statistical analysis

The experiments were designed taking into account good laboratory practices and the 3Rs principle of animal research. Both sexes of mice were used in all the experiments involving animal testing, except for the *in vivo* fluoxetine experiment presented in Fig. 3b (samples from only male mice were used due to a low number of female mice available in the cohort). Parametric tests were preferentially used to gain statistical power. Exceptions were made whenever the data presented lacked homoscedasticity or when variables were discrete, in which cases non-parametric tests were chosen. The statistical tests used in each particular experiment and statistical values are described in the legend of figures, and statistical values are provided in Table 1. Differences were considered statistically significant when p<0.05. Statistical analysis and plots were made in GraphPad Prism 6 software. Data are presented as mean± SEM. Detailed information on the statistical analysis of each experiment and number of samples/animals used is given below.

**Table 1.**
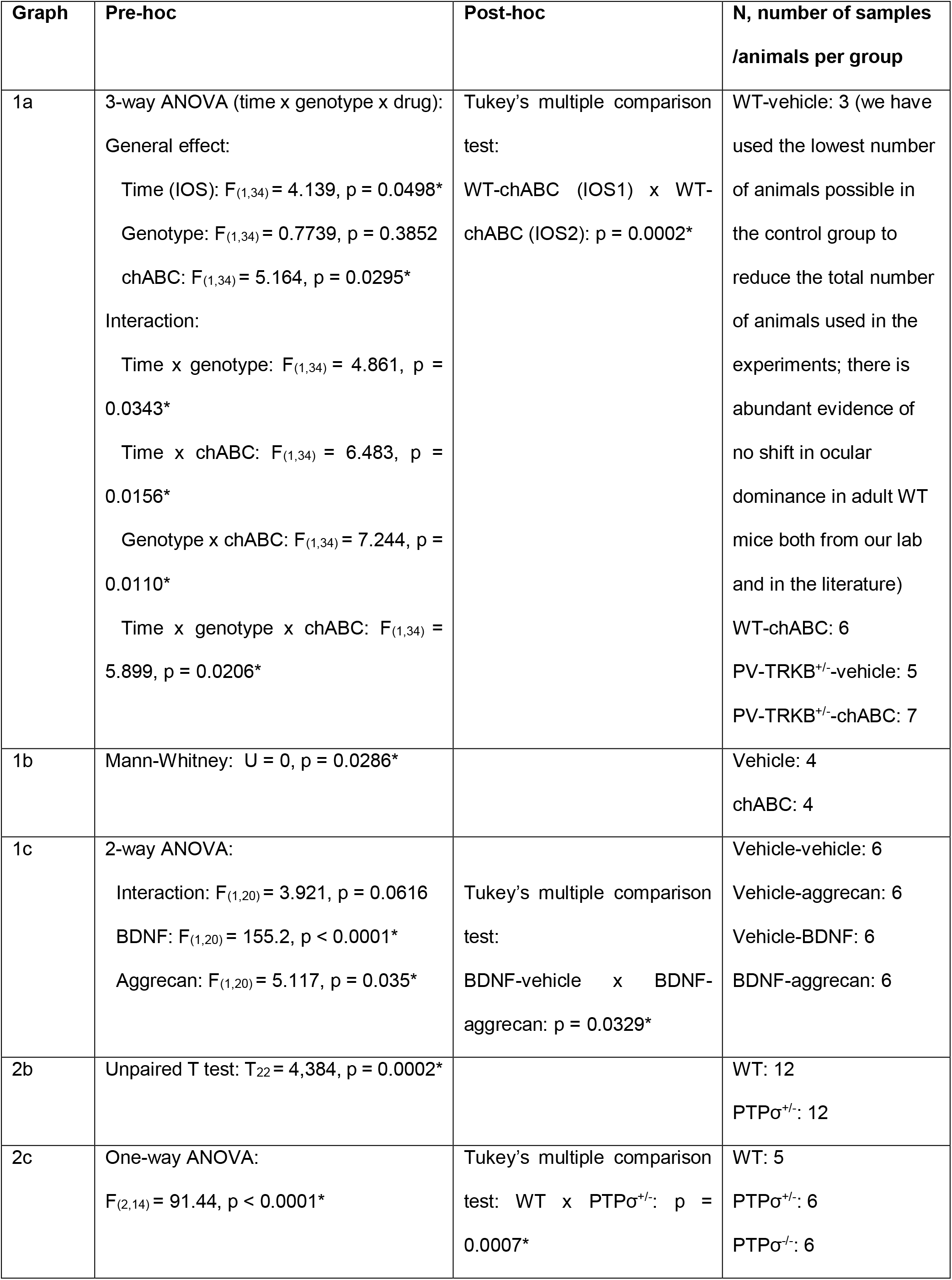
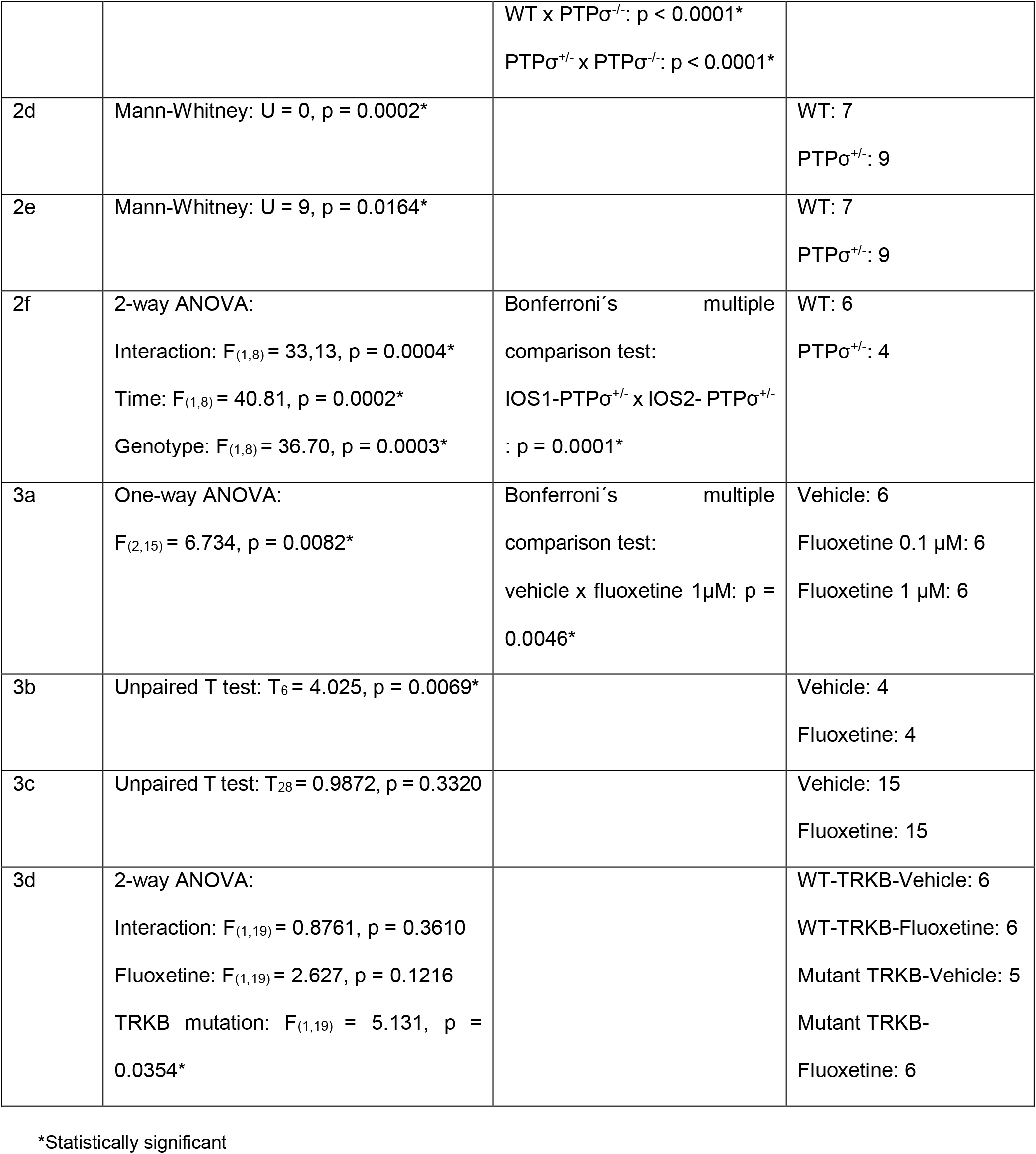
Statistical analysis and the number of animals/samples used in the experiments.

| Graph | Pre-hoc | Post-hoc | N, number of samples<br>/animals per group |
| --- | --- | --- | --- |
| 1a | 3-way ANOVA (time x genotype x drug):<br>General effect:<br>Time (IOS): $F_{(1,34)} = 4.139$, $p = 0.0498^*$<br>Genotype: $F_{(1,34)} = 0.7739$, $p = 0.3852$<br>chABC: $F_{(1,34)} = 5.164$, $p = 0.0295^*$<br>Interaction:<br>Time x genotype: $F_{(1,34)} = 4.861$, $p = 0.0343^*$<br>Time x chABC: $F_{(1,34)} = 6.483$, $p = 0.0156^*$<br>Genotype x chABC: $F_{(1,34)} = 7.244$, $p = 0.0110^*$<br>Time x genotype x chABC: $F_{(1,34)} = 5.899$, $p = 0.0206^*$ | Tukey's multiple comparison test:<br>WT-chABC (IOS1) x WT-chABC (IOS2): $p = 0.0002^*$ | WT-vehicle: 3 (we have used the lowest number of animals possible in the control group to reduce the total number of animals used in the experiments; there is abundant evidence of no shift in ocular dominance in adult WT mice both from our lab and in the literature)<br>WT-chABC: 6<br>PV-TRKB<sup>+/-</sup>-vehicle: 5<br>PV-TRKB<sup>+/-</sup>-chABC: 7 |
| 1b | Mann-Whitney: $U = 0$, $p = 0.0286^*$ | | Vehicle: 4<br>chABC: 4 |
| 1c | 2-way ANOVA:<br>Interaction: $F_{(1,20)} = 3.921$, $p = 0.0616$<br>BDNF: $F_{(1,20)} = 155.2$, $p < 0.0001^*$<br>Aggrecan: $F_{(1,20)} = 5.117$, $p = 0.035^*$ | Tukey's multiple comparison test:<br>BDNF-vehicle x BDNF-aggrecan: $p = 0.0329^*$ | Vehicle-vehicle: 6<br>Vehicle-aggrecan: 6<br>Vehicle-BDNF: 6<br>BDNF-aggrecan: 6 |
| 2b | Unpaired T test: $T_{22} = 4,384$, $p = 0.0002^*$ | | WT: 12<br>PTPσ<sup>+/-</sup>: 12 |
| 2c | One-way ANOVA:<br>$F_{(2,14)} = 91.44$, $p < 0.0001^*$ | Tukey's multiple comparison test: WT x PTPσ<sup>+/-</sup>: $p = 0.0007^*$ | WT: 5<br>PTPσ<sup>+/-</sup>: 6<br>PTPσ<sup>-/-</sup>: 6 |
|  |  | WT x PTPσ<sup>-/-</sup>: p < 0.0001*<br>PTPσ<sup>+/-</sup> x PTPσ<sup>-/-</sup>: p < 0.0001* |  |
| 2d | Mann-Whitney: U = 0, p = 0.0002* |  | WT: 7<br>PTPσ<sup>+/-</sup>: 9 |
| 2e | Mann-Whitney: U = 9, p = 0.0164* |  | WT: 7<br>PTPσ<sup>+/-</sup>: 9 |
| 2f | 2-way ANOVA:<br>Interaction: F<sub>(1,8)</sub> = 33,13, p = 0.0004*<br>Time: F<sub>(1,8)</sub> = 40.81, p = 0.0002*<br>Genotype: F<sub>(1,8)</sub> = 36.70, p = 0.0003* | Bonferroni's multiple<br>comparison test:<br>IOS1-PTPσ<sup>+/-</sup> x IOS2- PTPσ<sup>+/-</sup><br>: p = 0.0001* | WT: 6<br>PTPσ<sup>+/-</sup>: 4 |
| 3a | One-way ANOVA:<br>F<sub>(2,15)</sub> = 6.734, p = 0.0082* | Bonferroni's multiple<br>comparison test:<br>vehicle x fluoxetine 1μM: p =<br>0.0046* | Vehicle: 6<br>Fluoxetine 0.1 μM: 6<br>Fluoxetine 1 μM: 6 |
| 3b | Unpaired T test: T<sub>6</sub> = 4.025, p = 0.0069* |  | Vehicle: 4<br>Fluoxetine: 4 |
| 3c | Unpaired T test: T<sub>28</sub> = 0.9872, p = 0.3320 |  | Vehicle: 15<br>Fluoxetine: 15 |
| 3d | 2-way ANOVA:<br>Interaction: F<sub>(1,19)</sub> = 0.8761, p = 0.3610<br>Fluoxetine: F<sub>(1,19)</sub> = 2.627, p = 0.1216<br>TRKB mutation: F<sub>(1,19)</sub> = 5.131, p =<br>0.0354* |  | WT-TRKB-Vehicle: 6<br>WT-TRKB-Fluoxetine: 6<br>Mutant TRKB-Vehicle: 5<br>Mutant TRKB-Fluoxetine: 6 |

\*Statistically significant

## Results

### TRKB signaling in PV^+^ neurons is essential for chondroitinase ABC-induced plasticity

Digestion of PNNs by chABC is a well-known activator of plasticity (Fawcett et al., 2019), and it has been shown to induce critical period-like plasticity in the adult rodent visual cortex through degradation of CSPGs in the extracellular matrix (Pizzorusso et al., 2002). However, precise molecular mechanisms responsible for the plasticity-promoting effect of chABC remain largely unknown.

We tested the hypothesis that TRKB functioning in PV^+^ interneurons is involved in chABC-induced plastic changes. For that purpose, we generated mice heterozygous for full-length TRKB allele in PV^+^ interneurons (PV-TRKB^+/-^) (Minichiello et al., 1999). We assessed the ability of chABC to promote ocular dominance plasticity in their visual cortex. Four-month old mice underwent assessment of their ocular dominance by optical imaging of the intrinsic signal (IOS1) (Cang et al., 2005; Steinzeig et al., 2017). After the first imaging session, the mice were injected with chABC or vehicle into the binocular area of the visual cortex followed by monocular deprivation for 7 days. As expected, vehicle-treated wild-type mice failed to show any OD plasticity in the second imaging session (IOS2) right after the monocular deprivation, whereas wild-type mice treated with chABC showed a clear shift in the ocular dominance towards the non-deprived eye (Fig. 1a), as previously shown (Pizzorusso et al., 2002). However, PV-TRKB^+/-^ mice failed to show any shift in ocular dominance regardless of whether they were treated with chABC or not (Fig. 1a). These data demonstrate that chABC-mediated reactivation of visual cortical plasticity is dependent on intact TRKB signaling in the PV^+^ interneurons.

**Fig. 1:**
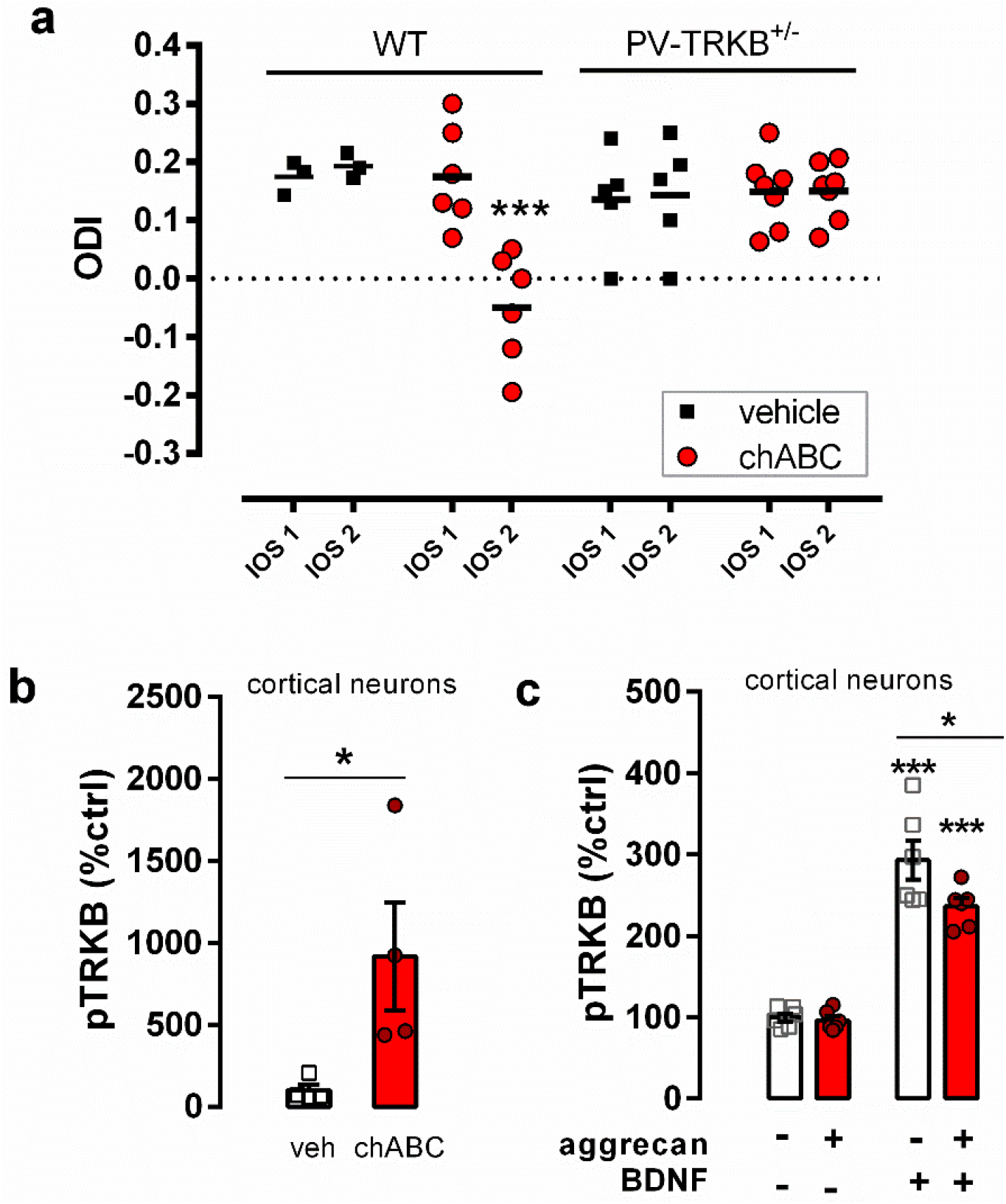
ChABC-induced plasticity requires intact TRKB signaling in PV^+^ neurons. a) PNN removal by local chABC injection reopens critical period-like plasticity in the visual cortex of WT but not in mice heterozygous for TRKB deletion in PV interneurons (PV-TRKB^+/-^), as measured by optical imaging of the intrinsic signal before (IOS1) and after 7 days of monocular deprivation (IOS2). ODI: ocular dominance index. b) ChABC treatment (30 min) of cortical neurons (7 DIV) increases TRKB phosphorylation (pTRKB). c) The CSPG aggrecan, added to cortical neurons (6 DIV) for 24 h, attenuates BDNF-induce TRKB phosphorylation in cortical neurons. Columns and bars represent mean±sem, respectively, and scattered points represent individual values. Data was analyzed by three-way ANOVA and Tukeys’s post-hoc (a), Mann-Whitney (b), or two-way ANOVA followed by Tukey’s multiple comparison test (c). *p<0.05, **p<0.005, ***p<0.0005.

Next, we assessed whether PNN digestion with chABC induces TRKB activation in primary cerebral cortical neurons. We treated rat cortical neurons grown in vitro (DIV) for 7 days with 2 U/ml chABC for 30 min and checked the activation of TRKB by assessing its phosphorylation levels (pTRKB) (Antila et al., 2014). We observed that reduction of PNNs by chABC treatment robustly increased pTRKB *in vitro* (Fig 1b).

After observing that PNN digestion with chABC treatment positively affects pTRKB, we investigated whether treatment with aggrecan, a major PNN component in the adult CNS, might have an opposing effect on pTRKB and render BDNF-induced TRKB activation less effective. We added 10 μg/ml aggrecan to the primary cortical neurons cultured for 6 DIV. After 24 hours, we challenged the cells with 20 ng/ml BDNF for 10 min. Aggrecan treatment did not alter basal pTRKB levels (that are normally very low *in vitro),* however, it significantly decreased BDNF-induced pTRKB (Fig. 1c). Taken together, these data demonstrate that PNNs regulate plasticity in a TRKB-dependent manner *in vivo* and that PNNs exert negative effects on TRKB activation *in vitro*.

### Deletion of CSPG receptor PTPσ extends the critical period and facilitates TRKB activation

PTPσ is a recognized inhibitor of neuronal plasticity and a receptor for CSPGs (Shen et al., 2009; Duan and Giger, 2010; Fawcett et al., 2019), and previous data indicate that PTPσ interacts with TRKB (Kurihara and Yamashita, 2012). We therefore hypothesized that PTPσ might restrict TRKB signaling through dephosphorylation in PNN-bearing PV^+^ interneurons and thereby inhibit TRKB-promoted plasticity. To confirm TRKB:PTPσ interaction, NIH3T3 cell line stably expressing either TRKA or TRKB (MG.TRKA and MG.TRKB, respectively), were transfected with myc-PTPσ, immunoprecipitated with anti-TRKB antibody and blotted for PTPσ. In the transfected samples, we observed a band below 250 kDa (Fig. 2a), which is consistent with the predicted molecular weight of the plasmid’s product. Additionally, a 165 kDa band was observed in non-transfected cells, which corresponds to the endogenous mature full-length PTPσ (Faux et al., 2007). When the membrane was stripped and reblotted for TRKB, we observed a band of 140 kDa corresponding to the full-length TRKB. This band was seen in TRKB expressing cells only, which rules out potential unspecific binding of TRKB antibodies to TRKA receptors. These data indicate that PTPσ interacts with TRKB. Furthermore, our previous proteomic study found PTPσ among the proteins that were immunoprecipitated with TRKB in hippocampal samples of the adult mouse brain (Fred et al., 2019).

**Fig. 2:**
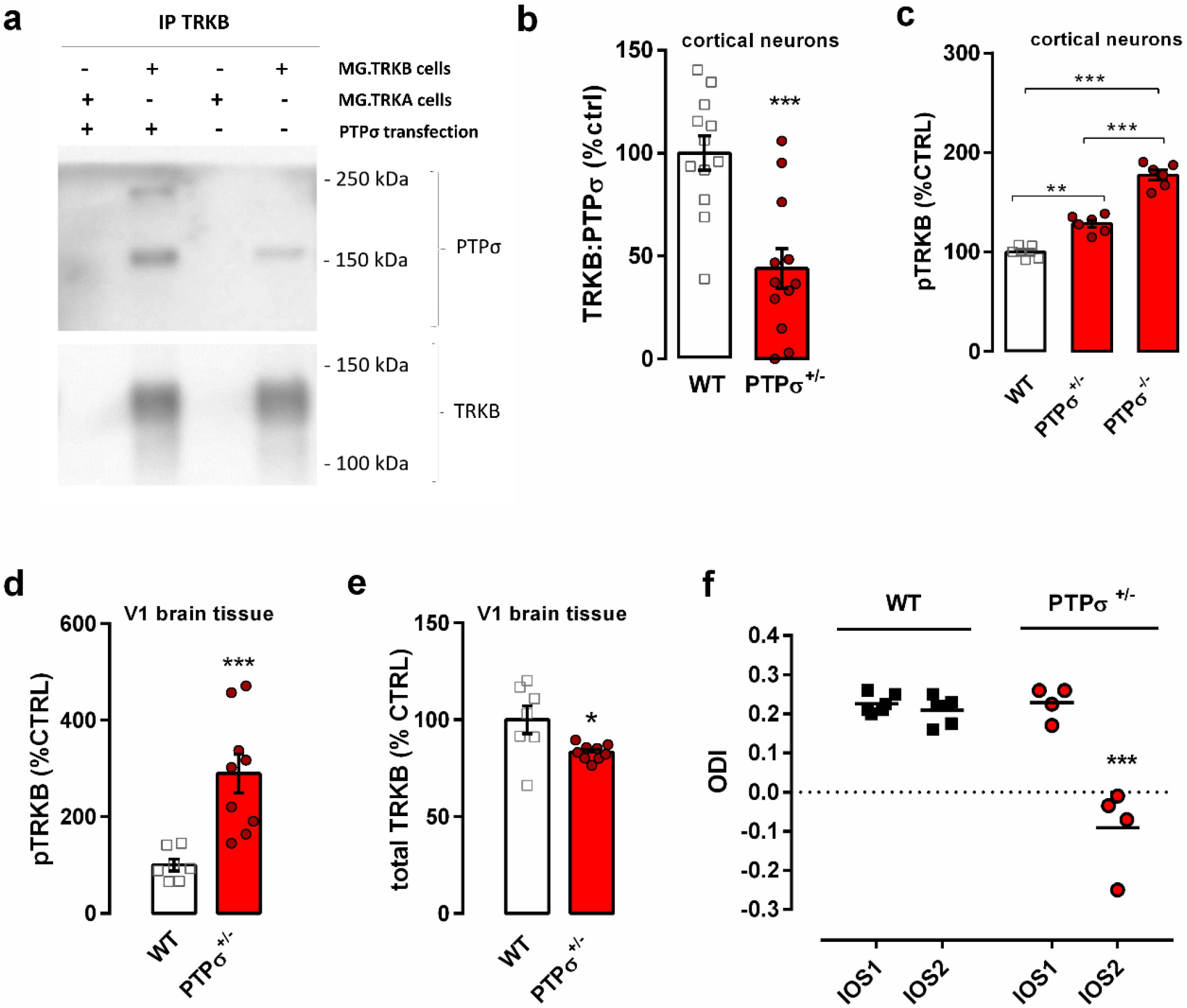
Deletion of CSPG receptor PTPσ facilitates TRKB phosphorylation and delays closure of critical period in the visual cortex of adult mice. a) PTPσ can be immunoprecipitated with anti-TRKB antibody in samples from TRKB expressing, but not from TRKA expressing cell line. b) Interaction of TRKB and PTPσ is reduced in embryonic cortical cultures from PTPσ+/-mice when compared to those from WT mice. c) TRKB phosphorylation is increased in cortical cultures from PTPσ^+/-^ and PTPσ^✓^’ mice. d) Adult PTPσ^+/-^ mice have increased TRKB phosphorylation and (e) a slight decrease in total TRKB levels in the visual cortex. f) Critical period-like plasticity is present in the visual cortex of two months old PTPσ^+/-^ mice (red circles) but not in WT littermates (black squares), as measured by optical imaging of the intrinsic signal before (IOS1) and after 3.5 days of monocular deprivation (IOS2). ODI: ocular dominance index. Columns and bars represent mean±sem, respectively, and scattered points represent individual values. Data was analyzed by two-way (f) or one-way ANOVA (c) followed by Bonferroni’s or Tukey’s post-hoc, respectively; unpaired T test (b) or Mann-Whitney test (d,e). *p<0.05, **p<0.005, ***p<0.0005.

Next, we set out to investigate the role of PTPσ in TRKB phosphorylation by using mice heterozygous for PTPσ (PTPσ^+/-^). Since transgenic mice quite often develop compensatory mechanisms to counteract genetic deficiency, we wanted to confirm that neurons lacking *PTPRS* allele would present comparable reduction in PTPσ:TRKB protein interaction. We prepared E18 neuronal cultures from cortex of WT or PTPσ^+/-^ littermates, and compared them after 7 DIV using co-immunoprecipitation. We observed a reduction of about 50% in the interaction between TRKB and PTPσ in samples from PTPσ^+/-^ mice as compared to WT (Fig. 2b). Then, we further investigated if PTPσ influences TRKB phosphorylation. We compared cortical pTRKB levels in embryonic cultures from PTPσ^-/-^ and PTPσ^+/-^ mice with those in cultures prepared from their WT littermates. Notably, PTPσ had a clear gene dosage-dependent suppressive effect on the basal pTRKB. PTPσ^+/-^ cortical cultures demonstrate increased pTRKB as compared to cultures from their WT littermates, and PTPσ^-/-^ samples demonstrate yet further increased pTRKB significantly differing from both PTPσ^+/-^ and WT samples (Fig. 2c). We also checked the levels of TRKB phosphorylation in samples from the adult visual cortex of mice heterozygous for *PTPRS* gene (PTPσ^+/-^). In line with our *in vitro* data, adult PTPσ^+/-^ mice exhibit increased basal TRKB autophosphorylation in the visual cortex (Fig. 2d), despite having slightly reduced levels of total TRKB (Fig. 2e), which may represent a compensatory downregulation due to a long-term increase in TRKB phosphorylation.

Finally, we investigated whether increased TRKB signaling produced by genetic deficiency of PTPσ might render the cortical structures of PTPσ^+/-^ mice susceptible to plastic changes. We tested adult PTPσ^+/-^ mice and their WT littermates in an ocular dominance plasticity paradigm using optical imaging. As expected, monocular deprivation did not induce an ocular dominance shift in adult WT animals; however, it did induce a shift in ocular dominance in PTPσ^+/-^ mice (Fig. 2f), indicating that closure of the critical period was delayed or prevented in the PTPσ^+/-^ mice.

### Antidepressant treatment disrupts TRKB:PTPσ interaction

Antidepressants have been demonstrated to induce activation of TRKB receptors in the brain (Saarelainen et al., 2003; Rantamäki et al., 2007; Castrén and Rantamäki, 2010). Moreover, fluoxetine, a widely used antidepressant, induces ocular dominance plasticity in the adult visual cortex through BDNF-TRKB signaling (Maya Vetencourt et al., 2008) and reduces percentage of PNNs enwrapping PV+ interneurons in the amygdala and hippocampus, shifting PV^+^ interneurons towards an immature state and reopening brain plasticity (Karpova et al., 2011). Therefore, we asked whether fluoxetine treatment might have an effect on TRKB: PTPσ interaction, which could potentially establish a link between antidepressant-induced TRKB phosphorylation and PNNs. We cultured rat primary cortical neurons for 7 DIV and challenged them with two different doses of fluoxetine (0.1 and 1 μM) for 30 min. Fluoxetine dose-dependently reduced the interaction between TRKB and PTPσ *in vitro* as measured by ELISA (Fig 3a). Since fluoxetine has been shown to increase TRKB phosphorylation as fast as 30 min after systemic injection (Saarelainen et al., 2003), we checked if fluoxetine would be able to disrupt TRKB: PTPσ interaction *in vivo* within a similar timeframe. We treated animals with 30 mg/kg fluoxetine (i.p.) and sacrificed the animals 30 minutes after the injection. Acute fluoxetine treatment significantly reduced TRKB:PTPσ interaction *in vivo* (Fig 3b).

**Fig. 3:**
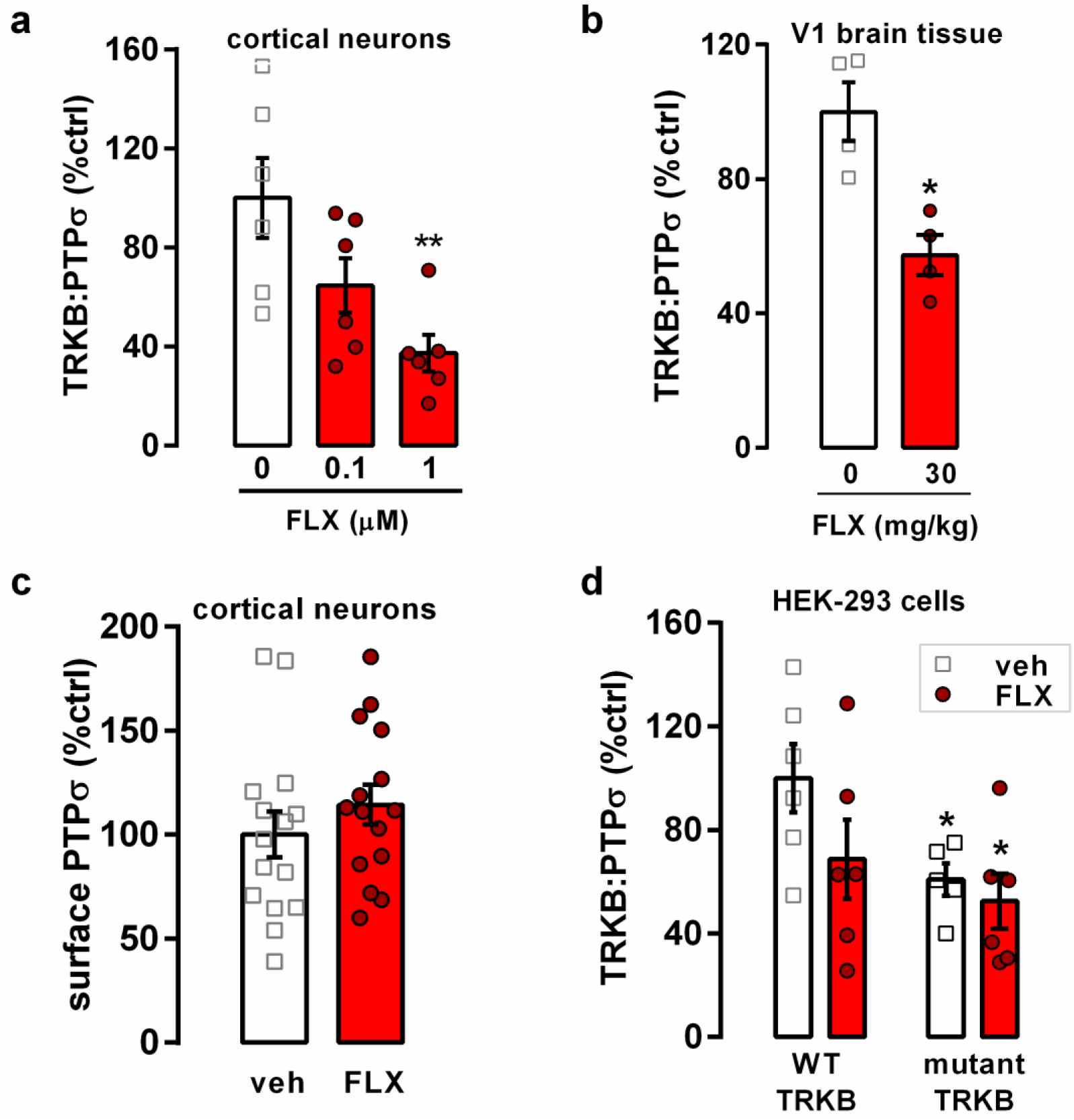
Fluoxetine disrupts TRKB and PTPσ interaction *in vitro* and *in vivo.* a) Fluoxetine (FLX) disrupts the interaction between TRKB and PTPσ in 7 DIV cortical neurons, and (b) in the visual cortex (V1) of mice systemically treated (30 mg/kg i.p.) 30 min before tissue dissection. c) Fluoxetine does not affect the detection of PTPσ extracellular domain on the cell surface, indicating no effect on PTPσ surface exposure or PTPσ ectodomain shedding. d) Mutation of the TRKB transmembrane domain (R427A/Y433F) partially disrupts the TRKB: PTPσ interaction and abolished the effects of fluoxetine on it. Columns and bars represent mean±sem, respectively. Data was analyzed by one-way ANOVA followed by Bonferroni’s (a), two-way ANOVA (d), or unpaired t test (b and c). *p<0.05, **p<0.005

PTPσ is proteolytically processed by cleavage in the juxtamembrane region, leading to shedding of the extracellular domain (Faux et al., 2007). To rule out the possibility that TRKB:PTPσ interaction experiments could have been influenced by a potential effect of the treatment upon PTPσ shedding, we investigated the presence of PTPσ extracellular domain in the cell surface after fluoxetine treatment in cortical cells using cell-surface ELISA (Zheng et al., 2008; Fred et al., 2019). We found no effect of the drug (Fig 3c). These data indicate that fluoxetine does not affect PTPσ shedding and positioning on cell surface, and supports the idea that the decrease in PTPσ levels in co-IP experiments is a result of decreased interaction with TRKB.

Antidepressants have recently been shown to directly interact with TRKB through the transmembrane region (Casarotto et al., 2019). Moreover, it has been previously suggested that PTPσ interacts with TRKA receptors through its transmembrane region (Faux et al., 2007). Therefore, we asked whether disruption in TRKB:PTPσ interaction after fluoxetine treatment is potentially mediated by the antidepressant’s binding to the same region of TRKB where interaction with PTPσ takes place. We transfected HEK293T cells with either a wild-type TRKB plasmid or a TRKB plasmid carrying two point mutations in the transmembrane region of TRKB (arginine R427 mutated to alanine, R427A; and tyrosine Y433 mutated to phenylalanine, Y433F); these point mutations disrupt a cholesterol interaction site in the TRKB transmembrane region critical for antidepressant interaction (Cannarozzo et al., 2020). R427A/Y433F mutation caused a dramatic decrease in TRKB: PTPσ interaction levels, however, the interaction was not completely lost, suggesting that multiple sites of interaction may exist. Nevertheless, fluoxetine (10 μM) failed to influence TRKB: PTPσ interaction in the cells carrying R427A/Y433F mutated TRKB (Fig. 3d), providing evidence that the site of interaction of PTPσ and fluoxetine with TRKB lies within the transmembrane region of TRKB.

## Discussion

In the current study, we have investigated a possible convergence of the two plasticity-inducing methods, chABC and antidepressant treatment, on the same molecular pathway involving reduced dephosphorylation of TRKB by PTPσ within PV^+^ interneurons. Perineuronal nets, extracellular structures rich in CSPGs, are associated with a reduction of plasticity in the adult brain. Maturation of PNNs coincides with the closure of the critical period (Berardi et al., 2004; Fawcett et al., 2019), and digestion of PNNs has been shown to promote plasticity and re-open critical periods for certain functions in the adult nervous system (Fawcett et al., 2019). Local injections of chondroitinase ABC promote axonal regeneration and functional recovery after spinal cord injury (Bradbury et al., 2002), restore ocular dominance plasticity in the visual cortex (Pizzorusso et al., 2002), and promote extinction of fear memories in rodents (Gogolla et al., 2009). Perineuronal nets preferentially enwrap PV^+^ interneurons that synchronize oscillatory activity of brain networks and mediate neuronal plasticity and learning. Our experiments demonstrate that the effects of chABC depend on TRKB signaling in PV^+^ interneurons, since genetic deficiency of TRKB in PV^+^ cells abrogates the ability of chondroitinase ABC to induce plasticity in the visual cortex of adult mice. Moreover, we have shown that chABC treatment increases pTRKB, which promotes TRKB signaling, while treatment with CSPG aggrecan decreases phosphorylation of TRKB induced by BDNF.

Chondroitin sulfate proteoglycans, major components of perineuronal nets, have been shown to exert inhibitory action on plasticity through protein tyrosine phosphatase receptor type S (Shen et al., 2009). PTPσ has been demonstrated to be critical for the inhibitory effects of CSPGs on plasticity and regeneration since CSPGs exert little inhibitory effect on axonal outgrowth of neurons prepared from PTPσ^✓^’ mice as compared to WT (Shen et al., 2009). Interestingly, PTPσ has been shown to interact with and dephosphorylate all three TRK receptors and inhibit their activation even in the presence of their cognate neurotrophins (Faux et al., 2007). Faux et al. provided evidence that PTPσ forms stable complexes with TRKA and TRKC and only weakly interacts with TRKB. However, subsequent research has shown that PTPσ co-immunoprecipitates with TRKB in samples from cortical neurons and mediates abrogation of BDNF-induced dendritic spine formation by CSPGs (Kurihara and Yamashita, 2012). We have now demonstrated that PTPσ dephosphorylates TRKB in *vitro* and in *vivo* based on increased phosphorylation of TRKB in cultures prepared from PTPσ KO homozygous or heterozygous mice (PTPσ^-/-^ and PTPσ^+/-^) and in samples from the visual cortex of PTPσ^+/-^ mice. Moreover, we observed that genetic deficiency of PTPσ delays the closure of critical period-like plasticity in the adult visual cortex, an effect that we demonstrated to be dependent on TRKB in PV^+^ neurons.

TRKB signaling is known to be critical for activation of plasticity by antidepressant drug treatment and for the behavioral consequences of it (Saarelainen et al., 2003; Duman and Monteggia, 2006; Autry and Monteggia, 2012; Castrén and Antila, 2017). Moreover, our group has recently demonstrated that TRKB activation in parvalbumin-positive interneurons is sufficient for the induction of juvenile-like plasticity in the adult brain and necessary for the plasticity-inducing effects of fluoxetine (Winkel et al., 2020). We now demonstrate that the antidepressant fluoxetine induces disruption of interaction between TRKB and PTPσ, releasing TRKB from the suppressive activity of the phosphatase and promoting its activation (Fig. 4). These data suggest that reduced dephosporylation of TRKB by PTPσ is at least one mechanism through which antidepressant treatment promotes plasticity.

**Figure 4.**
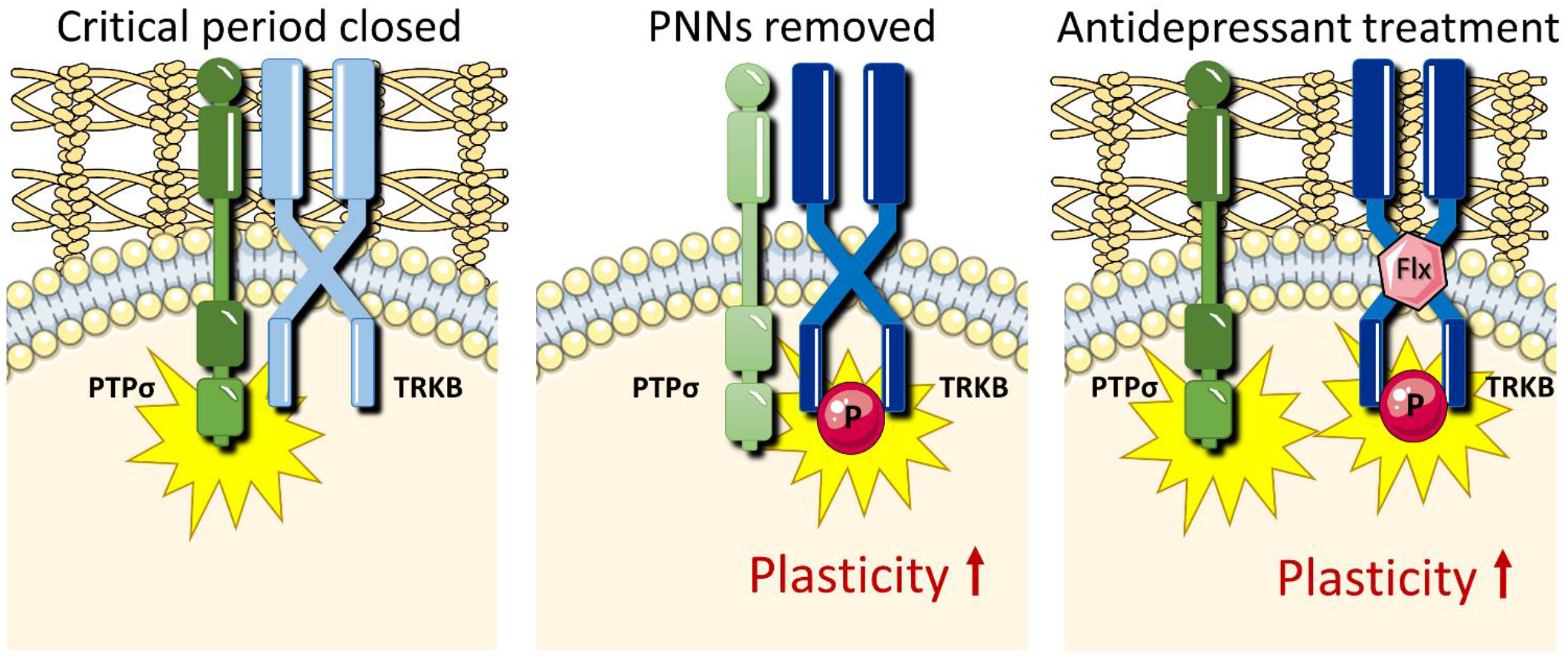
Schematic model of how chondroitinase and antidepressant treatments promote plasticity by releasing TRKB from dephosphorylating control of PTPσ. (Left): In the presence of PNNs, active PTPσ dephosphorylates TRKB and suppresses its signaling. (Middle): In the absence of PNNs, PTPσ is inactive and TRKB phosphorylation is facilitated. (Right): Fluoxetine (FLX) disrupts TRKB:PTPσ interaction, promoting TRKB signaling.

Antidepressants have recently been shown to directly interact with TRKB through its transmembrane domain (Casarotto et al., 2019). Our data now show that a mutation that inhibits antidepressant binding to the transmembrane region of TRKB partially disrupts its interaction with PTPσ, and that fluoxetine treatment has no further additive effect on TRKB: PTPσ interaction in TRKB mutant cells. These data suggest that TRKB and PTPσ interact in the transmembrane region of TRKB, and that binding of fluoxetine to TRKB disrupts TRKB: PTPσ interaction.

Our findings suggest a general mechanism responsible for the opening of critical period-like plasticity in the adult brain by chondroitinase ABC and antidepressant treatment (Fig. 4). We propose that both methods converge on the same pathway involving reduced inhibitory interaction between TRKB and PTPσ in PV^+^ neurons, which releases TRKB from the PTPσ-mediated dephosphorylation and promotes its autophosphorylation. Chondroitinase treatment downregulates CSPGs-mediated activation of PTPσ, which allows for enhanced signaling of TRKB (Fig. 4). Antidepressants, on the other hand, disrupt the interaction between TRKB and PTPσ in the membrane, promoting TRKB activation (Fig. 4).

Taken together, our data reveal that interaction between TRKB and PTPσ in PV^+^ interneurons is a critical regulator of chABC- and antidepressant-induced plasticity in the adult cortex. The similarity in the plasticity-promoting effects of chABC and antidepressants prompted us to focus on these two treatments, but there are a number of other methods known to reactivate juvenile-like plasticity in the adult cortex (Bavelier et al., 2010), such as enriched environment (Nithianantharajah and Hannan, 2006; Sale et al., 2007, 2014) and cross-modal manipulation of sensory functions (Rodríguez et al., 2018). It will be interesting in future studies to investigate whether induction of plasticity by these means recruits the same molecular pathways involving enhanced TRKB activation in PV^+^ interneurons.

## Author contributions

Conceptualization: CB and EC; Investigation: AL, PC, MF, MV, FW, AS, HA, JU, CB; Formal Analysis: AL, PC, MF, MV, AS, CB; Writing – Original Draft: AL, CB and EC; Writing – Review & Editing: AL, CB, EC, MF, MV, FW, AS, HA, PC, JU; Visualization: CB and AL; Supervision: CB and EC; Funding Acquisition: EC.

## Conflict of interest

The authors declare no conflict of interest.

## Acknowledgments

This work was supported by CIMO grant TM-17-10472 (to AL), the ERC grant #322742 - iPLASTICITY; EU Joint Programme - Neurodegenerative Disease Research (JPND) CircProt project #301225 and #643417, Doctoral Programme Brain & Mind, Sigrid Jusélius Foundation, Jane and Aatos Erkko Foundation, and the Academy of Finland grants #294710 and #307416 (to EC). The authors would like to thank Sulo Kolehmainen and Seija Lågas for technical help, Dr. Heikki Rauvala for the donation of the PTPσ knockout mice and for his expert comments on the manuscript, and Katja Kaurinkoski for language revision.

## Notes

### Competing Interest Statement

The authors have declared no competing interest.

### Summary of Updates

Some parts revised for clarity; graphical abstract revised and moved to Fig. 4

